# Serpine1 negatively regulates Th1 cell responses in experimental autoimmune encephalomyelitis

**DOI:** 10.1101/2023.07.13.548887

**Authors:** Irshad Akbar, Ruihan Tang, Joanie Baillargeon, Andrée-Pascale Roy, Prenitha Mercy Ignatius Arokia Doss, Chen Zhu, Vijay K. Kuchroo, Manu Rangachari

**Author notes:** Address correspondence and reprint request to Dr. Manu Rangachari or Dr. Vijay K. Kuchroo. Axe Neurosciences, Centre de recherche du CHU de Québec-Université Laval, 2705 boul Laurier, Québec QC G1V 4G2 (M.R) or Brigham and Women’s Hospital, Hale Building for Transformative Research, 60 Fenwood Rd, HBTM 10016F, Boston MA 02115. Email addresses. funding to I.A., MS Society of Canada Doctoral Studentship. funding to M.R., MS Society of Canada Discovery Grant #3781. funding to M.R., Canadian Institutes of Health Research (CIHR) Project Grant #159713. funding to M.R., Senior scholar award, Fonds de recherche de Québec – Santé #313330. Phone (M.R) : Fax (M.R).

## Abstract

Th1 cells are critical in experimental autoimmune encephalomyelitis (EAE). Serpine1 has been posited as an inhibitor of IFN_γ_ from T cells though its role in autoimmunity remains unclear. Here, we show that Serpine1 knockout (KO) mice develop EAE of enhanced severity relative to wild-type (WT) controls. Serpine1 overexpression represses Th1 cell cytokine production and pathogenicity, while Serpine1-KO:2D2 Th1 cells transfer EAE of increased severity in comparison to WT 2D2 Th1 cells. Notably, polarized Serpine1-KO Th1 cells display delayed expression of the Th1-specific inhibitory receptor, Tim-3. Serpine1-KO:Tim-3-Tg Th1 cells, which transgenically over-express Tim-3, showed increased expression of IFN_γ_ and reduced expression of the checkpoint molecules Lag-3 and PD-1 relative to WT Tim-3-Tg counterparts. Further, Serpine1 deficiency restored the EAE phenotype of Tim-3-Tg mice that normally develop mild disease. Together, we identify Serpine1 as a negative regulator of Th1 cells.

**Key points:**

- Serpine1 inhibits EAE in a T cell-dependent manner.
- Serpine1 is upregulated in Th1 cells and inhibits their pathogenicity.
- Serpine1 promotes expression and function of Th1-specific inhibitory receptor Tim-3.

## Introduction

Effector CD4^+^ Th1 cells are potent initiators and propagators of autoimmune diseases such as in the T cell-driven EAE^1^ model of MS^2^ (1). Th1 cells express a lineage-specific inhibitory receptor, Tim-3^3^, that resolves inflammatory responses when triggered at sites of inflammation (2–4). The Tim-3 pathway is crucial to repressing EAE pathology (2, 5, 6). Understanding the processes regulating Tim-3 expression and function might permit us to develop strategies to curb Th1-mediated inflammation in self-tissue.

Serpine1^4^, or PAI-1^5^, represses the conversion of the plasminogen proenzyme into mature plasmin and was previously suggested to regulate IFN_γ_-driven T cell responses (7, 8). Here, we show that Serpine1 restrains EAE pathogenicity via its inhibitory effects on Th1 cells, and that Serpine1 is required for optimal expression of Tim-3 by Th1 cells.

## Materials and Methods

### Mice

Tim-3-Tg (5) are described. B6^6^ WT^7^ (stock #000664), Serpine1-KO^8^ (B6.129S2-*Serpine1^tm1Mlg^*/J; #002507), 2D2-Tg (C57BL/6-Tg (Tcra2D2,Tcrb2D2)1KuchJ; #006912) and *Rag1^-/-^* (B6.129S7-*Rag1^tm1Mom^*/J; #002216) mice were obtained from Jackson Labs. Experimental and control animals were co-housed at the animal facilities of CRCHU de Québec-Université Laval or Brigham & Women’s Hospital. All procedures were authorized by the Animal Care Committee of Université Laval or the Institutional Animal Care and Use Committee of Harvard University.

### T helper cell differentiation

CD4^+^ T cells were enriched from B6 spleens using anti-CD4 Microbeads (Miltenyi), purified as CD4+CD62L^hi^ using a FACSAria (BD) high-speed cell sorter and cultured in supplemented T cell media as described (3). They were differentiated (9) for 2 days with plate bound anti-CD3 and anti-CD28 (2 μg mL^-1^ each; BioXcell) into Th1 cells – 10 ng mL^-1^ rmIL-12 (R&D Biosystems) plus anti-IL-4 (10 μg mL^-1^, BioXcell), or Th17 – rhTGFβ (3 ng mL^-1^, Miltenyi) + rmIL-6 (20 ng mL^-1^, Miltenyi) + anti-IFNγ (10 μg mL^-1^, BioXcell). Cells were then transferred to uncoated tissue culture plates and cultured for an additional 3 days, with rmIL-2 (10 ng mL^-1^, Miltenyi) added to Th1, and rmIL-23 (20 ng mL^-1^, R&D Biosystems) added to Th17. For multiple rounds of polarization, cells were collected at d5 and then restimulated for a subsequent 5-day period as above. Tiplaxtinin (25 μM, Tocris) or equivalent volume of DMSO were maintained for 5 days of culture where indicated. Serpine1 cDNA was cloned into pMIG^9^ vector and RV^10^ gene transduction of Th1 or Th17 cells was conducted using our described spin-infection protocol (10).

### EAE

Active immunization was induced by s.c. injection of MOG_[35-55]_^11^ (CHU de Québec) in incomplete Freund’s adjuvant (Difco) supplemented with 5 mg mL^-1^ *Mycobacterium tuberculosis* extract (Fisher). A dose of 25 μg MOG_[35-55]_ per mouse was used to compare WT to Serpine1-KO mice, while 100 μg per mouse was used in EAE experiments involving Tim-3-Tg mice. In Figure 1D, passive EAE was induced by first immunizing Serpine1-WT 2D2 and Serpine1-KO 2D2 mice with MOG_[35-55]_. Nine days later, splenocytes were collected and stimulated *ex vivo* for 48 hours with MOG_[35-55]_ (20 μg mL^-1^), rmIL-12 and rmIL-23. Next, 20x10^6^ splenocytes were injected i.p. into unimmunized B6 recipients. Adoptive transfer of WT 2D2, Serpine1-KO:2D2 or RV-transduced 2D2 Th1 cells entailed i.v. injection (2x10^6^) into *Rag1^-/-^* recipients at d5 of culture. In all EAE experiments, mice received 200 ng pertussis toxin (List Biological Labs) i.p. at d0 and d2. Mice were assessed for clinical symptoms daily as previously described (9).

**Figure 1.**
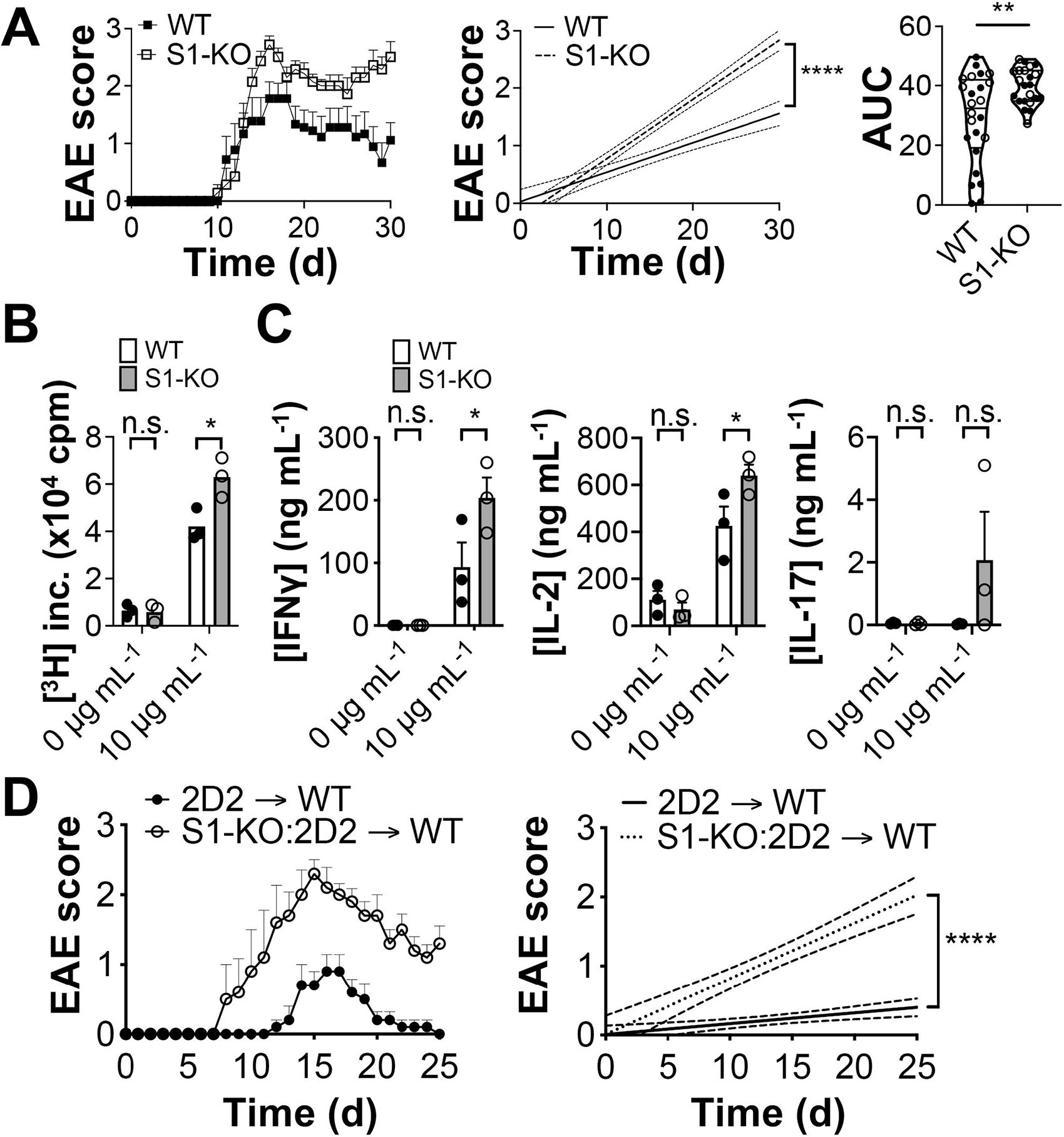
Serpine1 inhibits the severity of EAE. **A.** *Left,* Representative EAE curve of MOG_[35-55]_-immunized WT (n=9) and Serpine1-KO (n=7; abbreviated S1-KO) female mice. *Middle*, linear regression curves of the representative disease courses. The dashed lines indicate the 95% confidence intervals for each curve. *Right*, area-under-curve (AUC) comparison of mice pooled from 3 experiments. Filled circles, female mice (n=14, WT; n=12, Serpine1-KO); open circles, male mice (n=10, WT, n=10, Serpine1-KO). **, p<0.01; two way-ANOVA analysis of genotype as a variable. Incidence of EAE over all experiments was 24/25 WT, 22/24 Serpine1-KO. **B.** MOG_[35-55]_-immunized female WT and Serpine1-KO mice (n=3 each) were sacrificed 10 days post-immunization, and lymph node cells were stimulated, or not, with 10 µg mL^-1^ MOG_[35-55]_. Proliferation was assessed at 48 hours. cpm, counts per minute. **, p<0.01, *t*-test. **C.** Splenocytes from immunized female WT and Serpine1-KO mice (n=3 each) were restimulated, or not, with 10 µg mL^-1^ MOG_[35-55]_ for 48 (IL-2) or 72 (IFN_γ_, IL-17) hours, and secretion of the indicated cytokines was measured by ELISA. *, p<0.05; **, p<0.01, *t*-test. **D.** Splenocytes from female 2D2 and Serpine1-KO:2D2 mice were restimulated with MOG_[35-55]_ in the presence of IL-12 and IL-23, prior to adoptive transfer to WT mice (n=5 each condition) that were monitored for signs of EAE. Right graph, linear regression analysis with 95% confidence intervals. ****, p<0.0001.

### Flow cytometry

Cell surface and intracellular flow cytometry were conducted as previously described (10). The following Abs and dyes were used: *CD4*, clone RM4-5, ThermoFisher (TF) cats #45-0042-82, 48-0042-82; CD62L, MEL-14, TF #47-0621-82, *Tim-3*, RMT3-23, Biolegend #119706; *PD-1*, J43, TF #25-9985-82; *Lag-3*, eBioC9B7W, TF #12-2231-82; *IFNγ*, XMG1.2, TF # 48-7311-82; *TNFα*, MP6-XT22, TF #11-7321-82, 12-7321-41, 17-7321-82; IL-2, JES6-5H4, BD Biosciences #560547; IL-17, TC11-18H10.1, Biolegend #506922; Fixable Viability Dye, TF #65-0865-14; 7-aminoactinomycin D, TF #A1310. Data were collected using an LSRII flow cytometer (BD Biosciences) and were analyzed with FlowJo (BD). Gates were set on fluorescence minus one controls and the following global strategy was used: *i)* singlets were selected based on FSC-H vs FSC-A; *ii)* live CD4^+^ T cells based on CD4^+^ Viability Dye^neg^ events, or on CD4^+^7-aminoactinomycin D^neg^ events. RV-transduced CD4^+^ T cells were further gated on GFP positivity as indicated in Figure legends. In Figures 4CD, live CD4^+^ T cells were gated as CD4^+^Tim-3^pos^ or CD4^+^Tim-3^neg^.

### Ex vivo assessment of T cell function

Splenocyte cultures from EAE mice were stimulated, or not, with 10 μg mL^-1^ MOG_[35-55]_ for 48 hours. For proliferation studies, 1.25 μCi [^3^H]-thymidine (Perkin-Elmer) was added to each culture well for the last 16 hours. Cytokine supernatant ELISA were conducted using the following capture/detection sets: *IFNγ*, clones RA-6A2/XMG1.2; *IL-2*, JES6-1412/JES6-5H4; *IL-17*, TC11-18H10.1/TC11-8H4. Serpine1 protein was measured using Serpin E1/PAI-1 DuoSet ELISA (R&D Biosystems).

### Statistics

Two-tailed parametric tests were conducted using Prism (GraphPad). Comparisons of 2 groups were made by *t*-test while comparisons of >2 groups were made by ANOVA followed by post-hoc test. In EAE studies, linear regression (6, 10) or area under curve (10) analyses were conducted on mice with symptoms.

## Results and Discussion

### Serpine1 inhibits the severity of EAE

Upon immunization with MOG_[35-55]_, Serpine1-KO mice developed EAE of significantly greater severity than WT controls (Figure 1A). When the data were sex-disaggregated, Serpine1-KO females showed a significantly worsened disease burden as compared to WT females, while Serpine1-KO males showed a trend towards exacerbated disease relative to WT males (Supplementary Figure 1). Notably, antigen-specific proliferation responses to MOG_[35-55]_ were enhanced in peripheral Serpine1-KO CD4^+^ T cells prior to clinical onset (Figure 1B), as was secretion of IFN_γ_ and IL-2, but not IL-17 (Figure 1C).

Serpine1 regulates CNS fibrinolysis (11) and thus heightened EAE severity in Serpine1-KO mice might be due to its absence in a non-T cell compartment. We thus crossed Serpine1-KO mice to the 2D2-Tg strain, which bear a MOG_[35-55]_ -specific TCR (12), and immunized WT 2D2 and Serpine1-KO:2D2 mice. Prior to disease onset, we isolated splenocytes and restimulated them with Ag plus IL-12 and IL-23. Serpine1-KO:2D2 Ag-restimulated blasts induced EAE of significantly greater severity relative to WT 2D2 blasts upon adoptive transfer (Figure 1D).

Previous EAE studies that directly targeted Serpine1 *in vivo* revealed conflicting results possibly due to opposing effects on T cells and CNS repair (13, 14). Notably, in the chronic/relapsing Biozzi model of EAE, Serpine1-KO mice developed EAE of delayed onset and milder severity, due to superior CNS fibrinolytic capacity compared to controls (11); it is possible that the fibrogenic, pathogenic, properties of Serpine1 outweigh its anti-inflammatory function in this model. Here, we show that Serpine1 represses antigen-specific T cell-driven CNS autoimmunity in a T cell-intrinsic manner.

### Serpine1 suppresses Th1 cell function in vitro and in vivo

We next found that Serpine1 secretion was sharply upregulated in both Th1 (Figure 2A) and Th17 (Figure 2B) cells upon a second round of *in vitro* differentiation. Serpine1-deficient T cells can produce increased levels of IFN_γ_ relative *in vivo* (7, 8); however, the functional role of Serpine1 in differentiated *bonafide* Th1 and Th17 cells has never been directly assessed. Upon RV OE^12^ of Serpine1 in Th1 cells, we observed a downregulation in expression of IFN_γ_ and a trend towards reduced TNFα relative to control transduced cells. By contrast, overexpression of Serpine1 in Th17 cells did not impact expression of either IL-17 or TNFα (Figure 2C).

**Figure 2.**
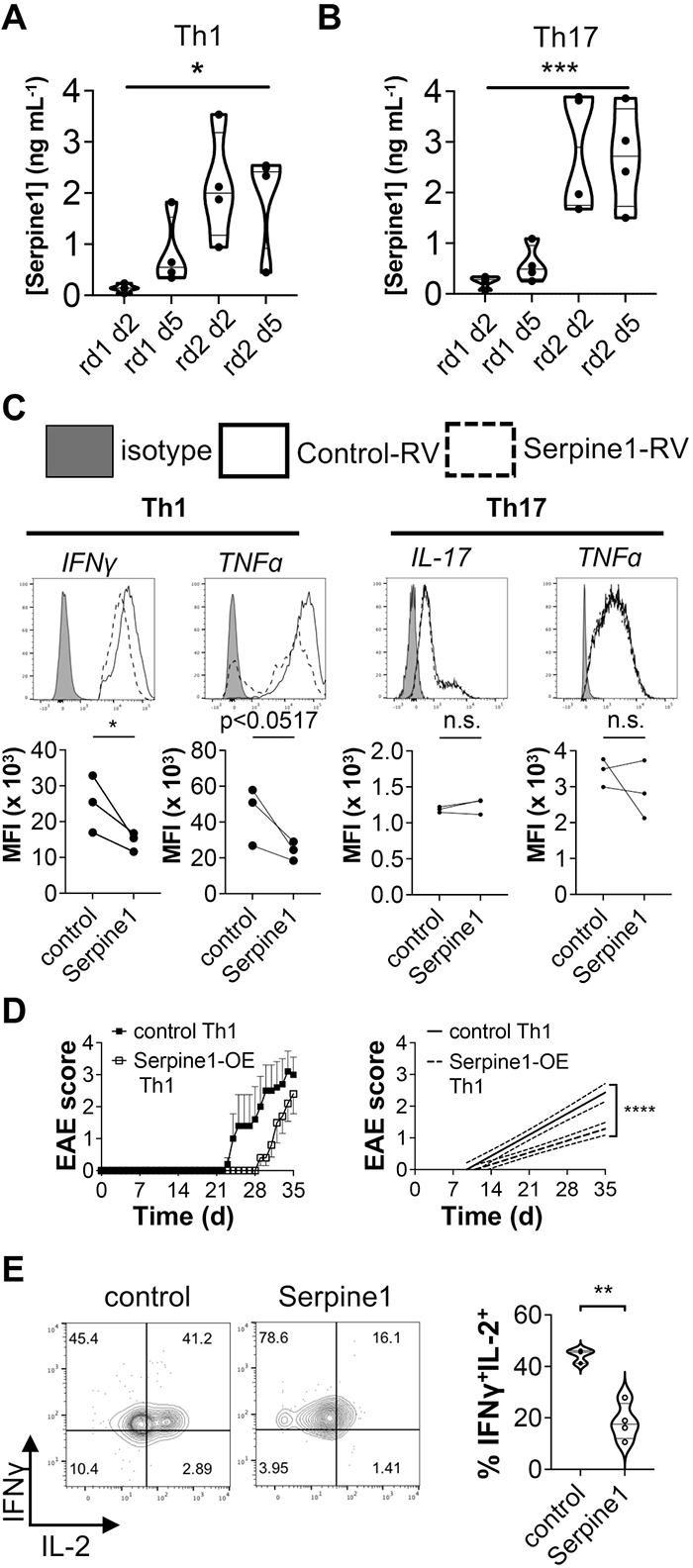
Serpine1 is expressed in Th1 cells and suppresses Th1-driven EAE. **A, B.** Naïve CD4^+^CD62L^hi^ T cells were isolated from female B6 mouse spleen and were differentiated under Th1 or Th17 conditions for 2 rounds of polarization. Secretion of Serpine1 protein was measured by ELISA in supernatant from Th1 (**A**) or Th17 (**B**) by ELISA. *, p<0.05; ***, p<0.001, one way ANOVA. Cultures derived from 4 independent mice. **C.** Female B6 Th1 or Th17 cells were transduced with control or Serpine1-OE RV. Cells were analyzed for production of the indicated cytokines after 5 days. Gated on GFP-positive live events. Quantitation represents paired *t*-test analysis of 3 independent cultures each. *, p<0.05. **D.** Female 2D2 Th1 cells were transduced with control-(n=5) or Serpine1-OE (n=4) RV. After 5 days of stimulation, cells were adoptively transferred to *Rag1^-/-^* recipients who were monitored for signs of EAE. *Bottom*, linear regression curves and 95% confidence intervals, for the disease courses. **E.** At disease endpoint (d35), splenic CD4^+^ T cells from mice in (**D**) were assessed for production of IFN_γ_ and IL-2 by flow cytometry. Gated on live CD4^+^ events. **, p<0.01, *t*-test.

To determine whether enforced expression of Serpine1 altered Th1 cell pathogenicity, we transduced 2D2 Th1 cells with Serpine1-OE or control RV and adoptively transferred these cells to *Rag1^-/-^*mice (6). Interestingly, Serpine1-OE 2D2 Th1 cells induced disease of lessened severity compared to control cells (Figure 2D), characterized by a reduced frequency of inflammatory IFNγ ^+^IL-2^+^ T cells *in vivo* (Figure 2E).

We next treated Th1 cells with tiplaxtinin, a small molecule that inhibits the antiproteolytic activity of Serpine1 towards the plasminogen activators urokinase and tissue plasminogen activator (15, 16). Tiplaxtinin enhanced IFN_γ_ and IL-2 production from Th1 cells (Figure 3A), suggesting that the Serpine1 may downregulate Th1 responses by repressing the activation of mature plasmin. We then generated Th1 cells from WT 2D2 and Serpine1-KO:2D2 mice and found that the latter induced EAE of significantly greater severity upon adoptive transfer (Figure 3B).

**Figure 3.**
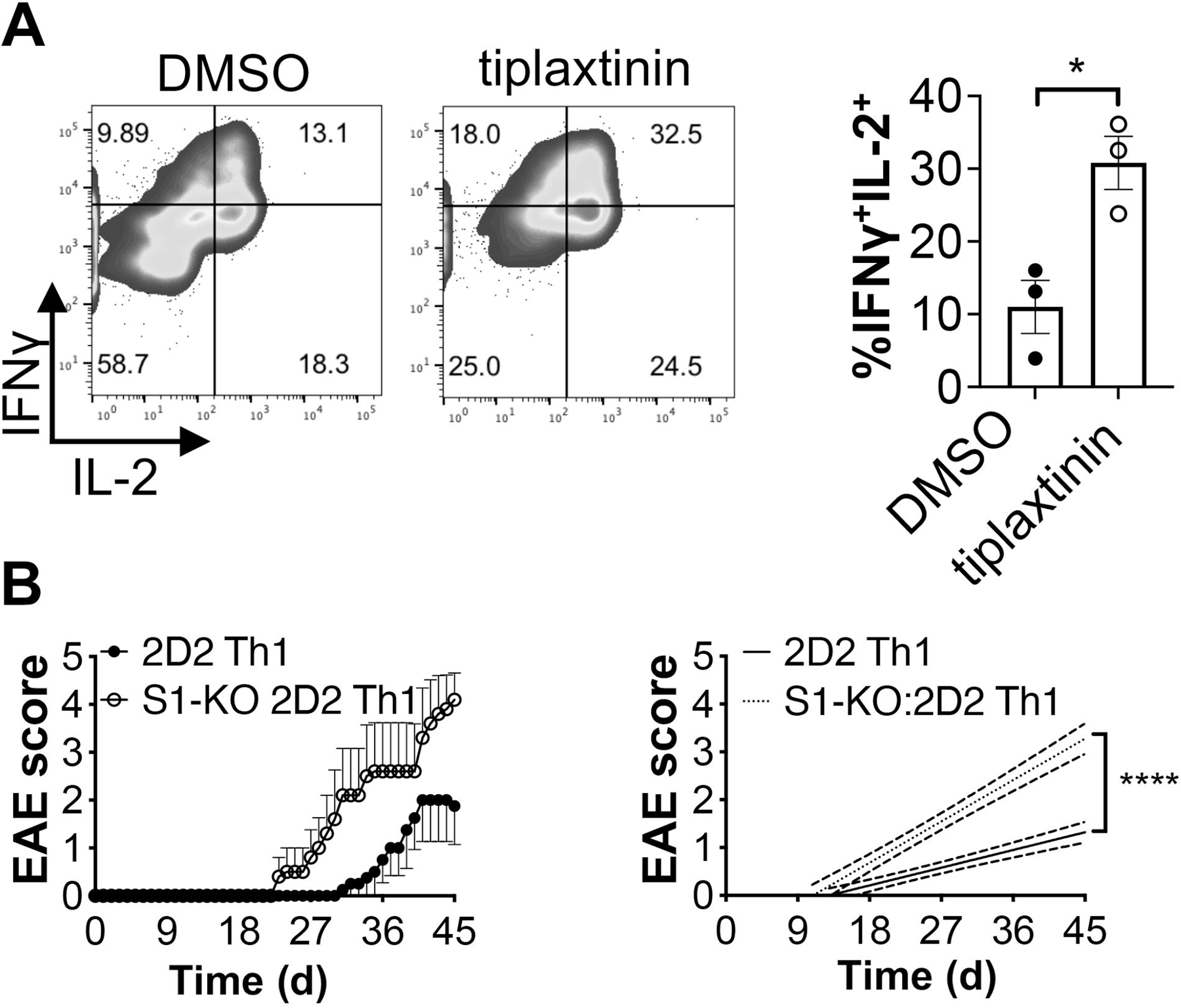
Loss of Serpine1 function or expression exacerbates Th1 responses. **A.** Female B6 Th1 cells were treated with DMSO (control) or 25 _μ_M tiplaxtinin for 5 days, at which point IFN_γ_ and IL-2 were measured by flow cytometry. *, p<0.05, t-test; 3 independent observations. **B.** Female 2D2 and Serpine1-KO:2D2 CD4^+^ Th1 cells were transferred i.v. to *Rag1^-/-^*recipients (2x10^6^ cells/mouse). n=4, 2D2; n=5, Serpine1-KO:2D2. Mice were subsequently assessed for signs of EAE. Left graph, linear regression curves of the disease courses.

Increased IFN_γ_ was previously observed from Serpine1-KO CD4^+^ and CD8^+^ T cells upon LPS or staphylococcal enterotoxin B treatment *in vivo* (7). Further, Serpine1-KO mice are resistant to nasal allergy in a Th2-mediated OVA sensitization model, with IFN_γ_ production upregulated by Serpine1-KO splenocytes upon Ag recall (8). Here, we show that while Serpine1 is expressed by both Th1 and Th17 cells, it impacts Th1 cells specifically by downregulating their inflammatory cytokine production and autoimmune potential.

### Serpine1 promotes Tim-3 expression and inhibits Tim-3^+^ Th1 cell inflammatory responses

As Serpine1 inhibits Th1-driven inflammation, we next asked whether it could augment the expression of inhibitory Tim-3. Repeatedly polarized Serpine1-KO Th1 cells expressed lower Tim-3, and with delayed kinetics, relative to S1-WT controls (Figure 4A). Loss of Serpine1 signaling did not increase expression of IFNγ; however, the frequency of IFNγ^+^Tim-3^+^ Th1 cells was strikingly lower in its absence (Figure 4B).

**Figure 4.**
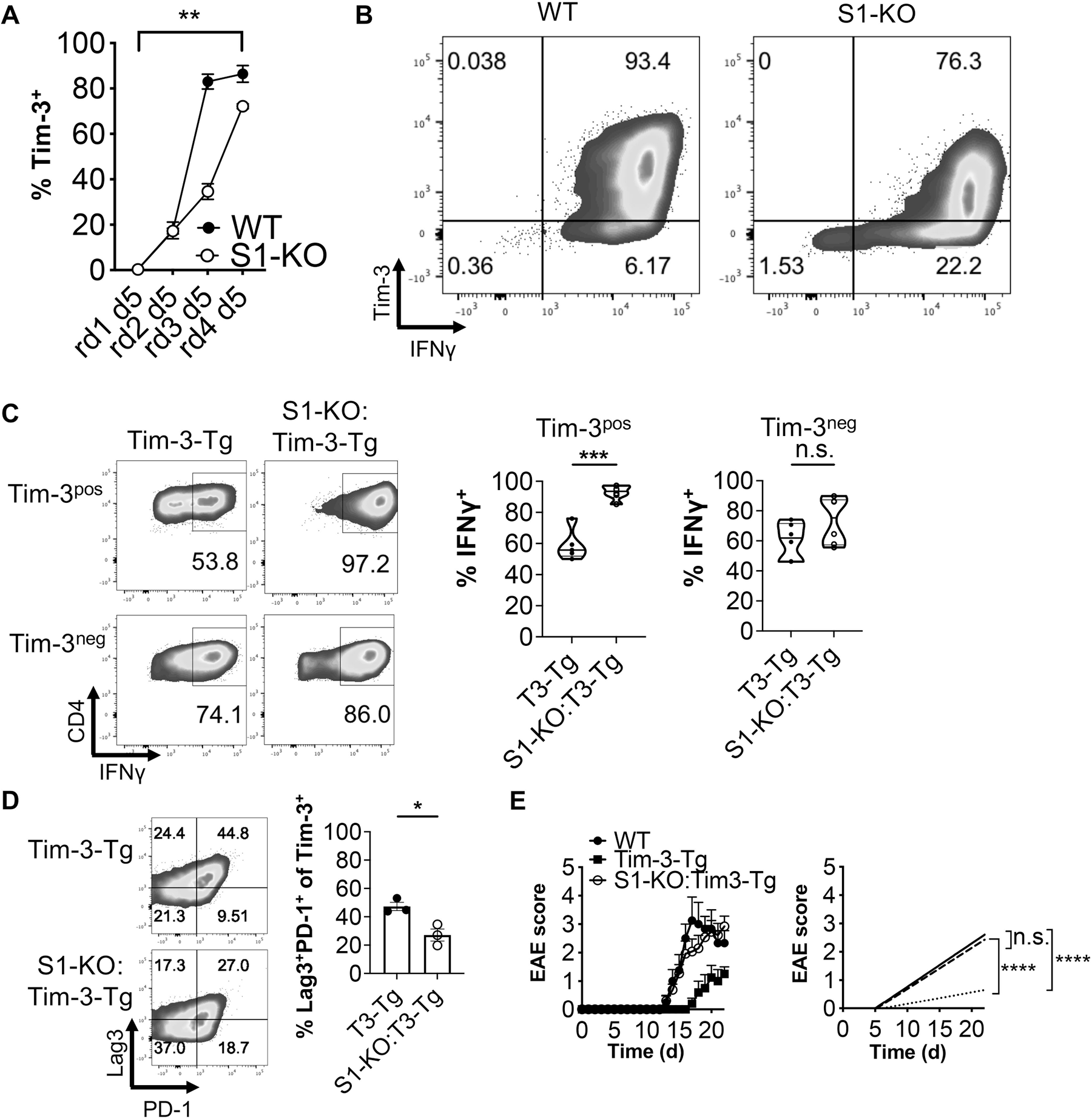
Serpine1 promotes Tim-3 expression and inhibits Tim-3^+^ Th1 cell inflammatory responses. **A, B.** Female WT and Serpine1-KO:T cells were subjected to 4 successive rounds of Th1 polarization, and Tim-3 (**A,B**) and IFN_γ_ (**B**) were assessed by flow cytometry. **(A)** Quantification of Tim-3 expression at the end of each round. **, p<0.01, one-way ANOVA. **B.** Expression of Tim-3 versus IFN_γ_ after 20 days of total culture (end of round 4). Gated on live CD4^+^ events. Representative of 4 experiments. **C.** Splenic T CD4^+^ T cells from male Tim-3-Tg and Serpine1-KO:Tim-3-Tg mice (n=6 independent cultures from each) were differentiated under Th1 conditions for 5 days. CD4, Tim-3 and IFNγ expression were assessed by flow cytometry, with cells first gated on live CD4^+^Tim-3^pos^ or live CD4^+^Tim-3^neg^. ***, p<0.001; *t*-test. **D.** Splenic T CD4^+^ T cells from male Tim-3-Tg and Serpine1-KO:Tim-3-Tg mice (n=3 independent cultures from each) were differentiated under Th1 conditions for 5 days. Lag-3 and PD-1 expression were assessed by flow cytometry. *, p<0.05; *t*-test. Gated on live CD4^+^Tim-3^+^ events. **E.** Female WT (n=4), WT Tim-3-Tg (n=5) and Serpine1-KO:Tim-3-Tg (n=14) and mice were immunized with MOG_[35-55]_ and were monitored for signs of EAE. *Right graph*, linear regression analysis of the disease curves with Bonferroni’s correction applied. Representative of 3 immunizations.

Tim-3-Tg mice ectopically overexpress Tim-3 cDNA in a T cell-restricted manner, without the need for multiple rounds of polarization under Th1 conditions (17). Reasoning that this might help us uncover Serpine1-dependent differences in IFNγ expression, we crossed Serpine1-KO mice to the Tim-3-Tg strain and generated Th1 cells from these and Tim-3-Tg control mice. Tim-3^pos^ Th1 cells derived from Serpine1-KO:Tim-3-Tg mice were strikingly more positive for IFNγ than Tim-3-Tg counterparts; notably, no differences were observed with Tim-3^neg^ Th1 cells between the strains (Figure 4C). This indicated that Serpine1 represses IFNγ expression in a Tim-3-dependent manner. Interestingly, there was a concomitant reduction in the expression of the T cell negative regulatory receptors PD-1 and Lag-3 in Serpine1-KO:Tim-3-Tg Th1 cells when compared to Tim-3-Tg controls (Figure 4D).

Tim-3-Tg mice develop EAE of attenuated severity (5). To examine whether loss of Serpine1 expression could reverse this phenotype, we actively immunized WT, Tim-3-Tg and Serpine1:KO Tim-3-Tg mice. While Tim-3-Tg mice developed disease of only mild severity as expected, Serpine1-KO:Tim-3-Tg mice displayed EAE that was of comparable severity to that seen in WT animals (Fig 4E). Our data thus demonstrate that Serpine1 is required for Tim-3-mediated repression of T cell inflammation and pathogenicity in EAE.

Tim-3 marks exhausted T cells in chronic viral infections and cancers (18–20) and is functionally tractable, as concomitant blockade of Tim-3 and PD-1 causes tumor regression greater degree than that with anti-PD-1 alone (20). Further, depletion of Bat3, an intracellular repressor of Tim-3 signaling, causes Th1 cells to adopt an exhausted-like phenotype *in vivo* (6, 21). While Serpine1-mediated upregulation of Tim-3 may be desirable in the context of autoimmunity, it remains to be seen what role, if any, Serpine1 plays in T cell exhaustion.

Altogether, our data identify a novel Th1 cell-intrinsic regulatory mechanism. Serpine1 represses Th1 cell pathogenicity by inhibiting their production of inflammatory cytokines and by restraining them from adopting a highly differentiated Tim-3^+^ phenotype. Strategies to augment Serpine1 expression and function in Th1 cells could present an attractive therapeutic option in autoimmune disease.

## Acknowledgements

We thank Vincent Desrosiers for technical assistance, Kim Larose-Labrecque and Andrée Brisson for animal care, and Ryder Whittaker Hawkins for critical reading of the manuscript.

## Author Contributions^13^

## Data availability statement^14^

## Footnotes

1 Experimental autoimmune encephalomyelitis

2 Multiple sclerosis

3 T cell immunoglobulin and mucin domain-containing-3

4 Serine protease inhibitor clade E1

5 plasminogen activation inhibitor 1

6 C57BL6/J

7 Serpine1-WT

8 Serpine1-KO

9 pMSCV-IRES-GFP

10 retroviral

11 Myelin oligodendrocyte glycoprotein, amino acids 35-55

12 overexpression

13 I.A. conducted experiments and managed the project. R.T., J.B., A-P.R., P.M.I.A.D and C.Z. conducted experiments. V.K.K. co-supervised the project. M.R. conducted experiments, supervised the project and wrote the manuscript.

14 The datasets generated for this study are available from the corresponding authors upon reasonable request.

15 V.K.K. has an ownership stake and is a member of the Scientific Advisory Board for Tizona Therapeutics. Further, he is a co-founder of, and has an ownership stake in, Celsius Therapeutics. In addition, he is an inventor on patents related to Th17 cell function. His interests are reviewed and managed by Brigham & Women’s Hospital and Partners Healthcare in accordance with their conflict of interest policies.

## References

1. Jäger, A., V. Dardalhon, R. A. Sobel, E. Bettelli, and V. K. Kuchroo. 2009. Th1, Th17, and Th9 effector cells induce experimental autoimmune encephalomyelitis with different pathological phenotypes. J. Immunol. 183: 7169–7177.

2. Monney, L., C. A. Sabatos, J. L. Gaglia, A. Ryu, H. Waldner, T. Chernova, S. Manning, E. A. Greenfield, A. J. Coyle, R. A. Sobel, G. J. Freeman, and V. K. Kuchroo. 2002. Th1-specific cell surface protein Tim-3 regulates macrophage activation and severity of an autoimmune disease. Nature 415: 536–541.

3. Sabatos, C. A., S. Chakravarti, E. Cha, A. Schubart, A. Sánchez-Fueyo, X. X. Zheng, A. J. Coyle, T. B. Strom, G. J. Freeman, and V. K. Kuchroo. 2003. Interaction of Tim-3 and Tim-3 ligand regulates T helper type 1 responses and induction of peripheral tolerance. Nat. Immunol. 4: 1102–1110.

4. Sánchez-Fueyo, A., J. Tian, D. Picarella, C. Domenig, X. X. Zheng, C. A. Sabatos, N. Manlongat, O. Bender, T. Kamradt, V. K. Kuchroo, J.-C. Gutiérrez-Ramos, A. J. Coyle, and T. B. Strom. 2003. Tim-3 inhibits T helper type 1-mediated auto- and alloimmune responses and promotes immunological tolerance. Nat. Immunol. 4: 1093–1101.

5. Dardalhon, V., A. C. Anderson, J. Karman, L. Apetoh, R. Chandwaskar, D. H. Lee, M. Cornejo, N. Nishi, A. Yamauchi, F. J. Quintana, R. A. Sobel, M. Hirashima, and V. K. Kuchroo. 2010. Tim-3/galectin-9 pathway: regulation of Th1 immunity through promotion of CD11b+Ly-6G+ myeloid cells. J. Immunol. 185: 1383–1392.

6. Rangachari, M., C. Zhu, K. Sakuishi, S. Xiao, J. Karman, A. Chen, M. Angin, A. Wakeham, E. A. Greenfield, R. A. Sobel, H. Okada, P. J. McKinnon, T. W. Mak, M. M. Addo, A. C. Anderson, and V. K. Kuchroo. 2012. Bat3 promotes T cell responses and autoimmunity by repressing Tim-3–mediated cell death and exhaustion. Nat. Med. 18: 1394–1400.

7. Renckens, R., J. M. Pater, and T. van der Poll. 2006. Plasminogen activator inhibitor type-1-deficient mice have an enhanced IFN-gamma response to lipopolysaccharide and staphylococcal enterotoxin B. J. Immunol. 177: 8171–8176.

8. Sejima, T., S. Madoiwa, J. Mimuro, T. Sugo, K. Okada, S. Ueshima, O. Matsuo, T. Ishida, K. Ichimura, and Y. Sakata. 2005. Protection of plasminogen activator inhibitor-1-deficient mice from nasal allergy. J. Immunol. 174: 8135–8143.

9. Pradeep Yeola, A., I. Akbar, J. Baillargeon, P. Mercy Ignatius Arokia Doss, V. O. Paavilainen, and M. Rangachari. 2020. Protein translocation and retro-translocation across the endoplasmic reticulum are crucial to inflammatory effector CD4+ T cell function. Cytokine 129: 154944.

10. Doss, P. M. I. A., M. Umair, J. Baillargeon, R. Fazazi, N. Fudge, I. Akbar, A. P. Yeola, J. B. Williams, M. Leclercq, C. Joly Beauparlant, P. Beauchemin, G. F. Ruda, M. Alpaugh, A. C. Anderson, P. E. Brennan, A. Droit, H. Lassmann, C. S. Moore, and M. Rangachari. 2021. Male sex chromosomal complement exacerbates the pathogenicity of Th17 cells in a chronic model of central nervous system autoimmunity. Cell Rep 34: 108833.

11. East, E., D. Gveric, D. Baker, G. Pryce, H. R. Lijnen, and M. L. Cuzner. 2008. Chronic relapsing experimental allergic encephalomyelitis (CREAE) in plasminogen activator inhibitor-1 knockout mice: the effect of fibrinolysis during neuroinflammation. Neuropathol. Appl. Neurobiol. 34: 216–230.

12. Bettelli, E., M. Pagany, H. L. Weiner, C. Linington, R. A. Sobel, and V. K. Kuchroo. 2003. Myelin oligodendrocyte glycoprotein-specific T cell receptor transgenic mice develop spontaneous autoimmune optic neuritis. J. Exp. Med. 197: 1073–1081.

13. Gur-Wahnon, D., T. Mizrachi, F.-Y. Maaravi-Pinto, A. Lourbopoulos, N. Grigoriadis, A.-A. R. Higazi, and T. Brenner. 2013. The plasminogen activator system: involvement in central nervous system inflammation and a potential site for therapeutic intervention. J Neuroinflammation 10: 124.

14. Pelisch, N., T. Dan, A. Ichimura, H. Sekiguchi, D. E. Vaughan, C. van Ypersele de Strihou, and T. Miyata. 2015. Plasminogen Activator Inhibitor-1 Antagonist TM5484 Attenuates Demyelination and Axonal Degeneration in a Mice Model of Multiple Sclerosis. PLoS ONE 10: e0124510.

15. Elokdah, H., M. Abou-Gharbia, J. K. Hennan, G. McFarlane, C. P. Mugford, G. Krishnamurthy, and D. L. Crandall. 2004. Tiplaxtinin, a novel, orally efficacious inhibitor of plasminogen activator inhibitor-1: design, synthesis, and preclinical characterization. J. Med. Chem. 47: 3491–3494.

16. Gorlatova, N. V., J. M. Cale, H. Elokdah, D. Li, K. Fan, M. Warnock, D. L. Crandall, and D. A. Lawrence. 2007. Mechanism of inactivation of plasminogen activator inhibitor-1 by a small molecule inhibitor. J. Biol. Chem. 282: 9288–9296.

17. Dardalhon, V., A. S. Schubart, J. Reddy, J. H. Meyers, L. Monney, C. A. Sabatos, R. Ahuja, K. Nguyen, G. J. Freeman, E. A. Greenfield, R. A. Sobel, and V. K. Kuchroo. 2005. CD226 is specifically expressed on the surface of Th1 cells and regulates their expansion and effector functions. J. Immunol. 175: 1558–1565.

18. Jones, R. B., L. C. Ndhlovu, J. D. Barbour, P. M. Sheth, A. R. Jha, B. R. Long, J. C. Wong, M. Satkunarajah, M. Schweneker, J. M. Chapman, G. Gyenes, B. Vali, M. D. Hyrcza, F. Y. Yue, C. Kovacs, A. Sassi, M. Loutfy, R. Halpenny, D. Persad, G. Spotts, F. M. Hecht, T.-W. Chun, J. M. McCune, R. Kaul, J. M. Rini, D. F. Nixon, and M. A. Ostrowski. 2008. Tim-3 expression defines a novel population of dysfunctional T cells with highly elevated frequencies in progressive HIV-1 infection. J. Exp. Med. 205: 2763–2779.

19. Jin, H.-T., A. C. Anderson, W. G. Tan, E. E. West, S.-J. Ha, K. Araki, G. J. Freeman, V. K. Kuchroo, and R. Ahmed. 2010. Cooperation of Tim-3 and PD-1 in CD8 T-cell exhaustion during chronic viral infection. Proc. Natl. Acad. Sci. U.S.A. 107: 14733–14738.

20. Sakuishi, K., L. Apetoh, J. M. Sullivan, B. R. Blazar, V. K. Kuchroo, and A. C. Anderson. 2010. Targeting Tim-3 and PD-1 pathways to reverse T cell exhaustion and restore anti-tumor immunity. J. Exp. Med. 207: 2187–2194.

21. Zhu, C., K. O. Dixon, K. Newcomer, G. Gu, S. Xiao, S. Zaghouani, M. A. Schramm, C. Wang, H. Zhang, K. Goto, E. Christian, M. Rangachari, O. Rosenblatt-Rosen, H. Okada, T. Mak, M. Singer, A. Regev, and V. Kuchroo. 2021. Tim-3 adaptor protein Bat3 is a molecular checkpoint of T cell terminal differentiation and exhaustion. Sci Adv 7.

