## Supplemental Figure 1 for "Serpine1 negatively regulates Th1 cell responses in experimental autoimmune encephalomyelitis"

### Slide 1
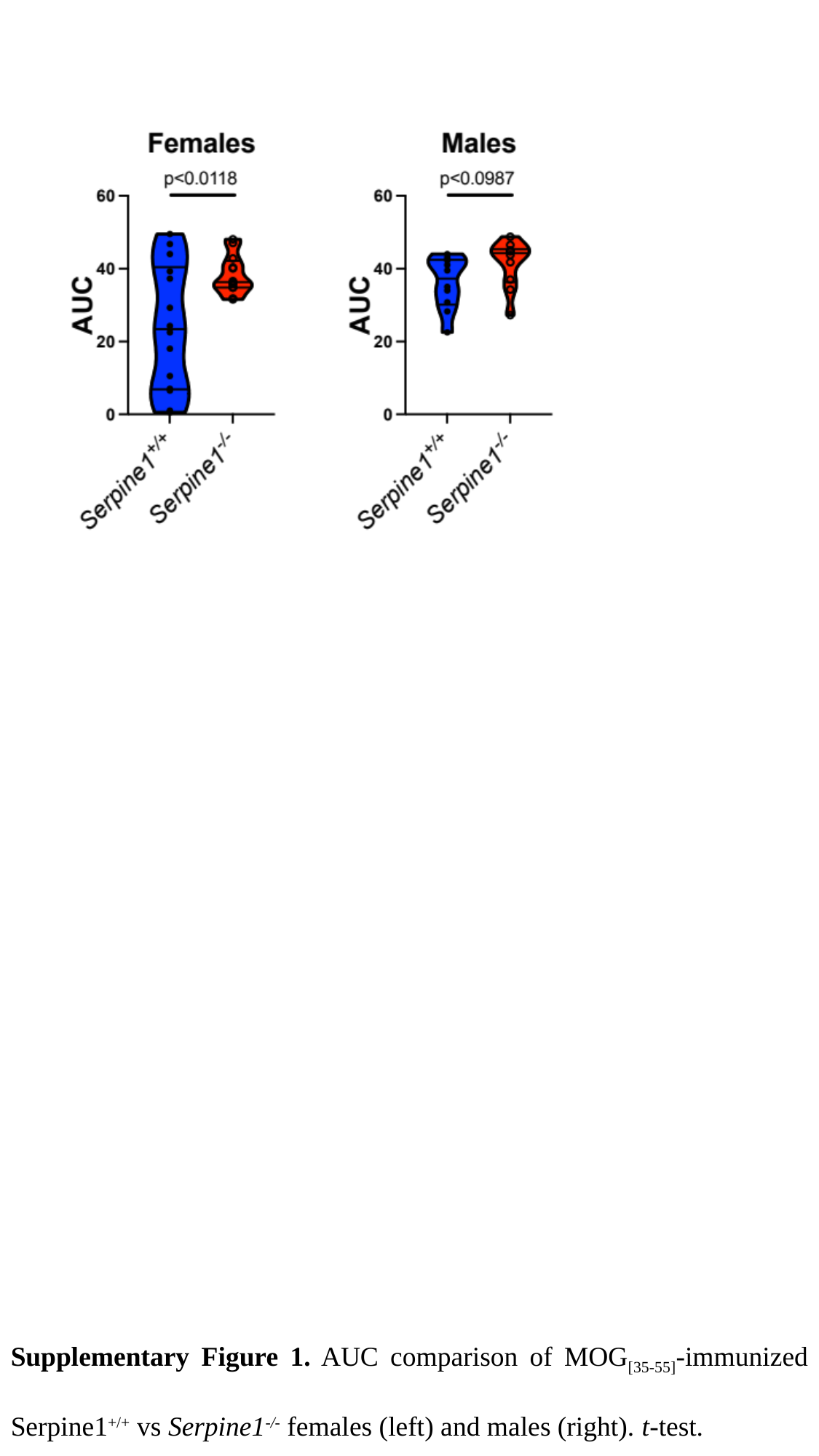

Supplementary Figure 1. AUC comparison of MOG_[35-55]_-immunized Serpine1^+/+^ vs Serpine1^-/-^ females (left) and males (right). t-test.
